# Establishment of a transgenic strain for the whole brain calcium imaging in larval medaka fish (*Oryzias latipes*)

**DOI:** 10.1101/2025.04.09.647916

**Authors:** Takahide Seki, Kazuhiro Miyanari, Asuka Shiraishi, Sachiko Tsuda, Satoshi Ansai, Hideaki Takeuchi

## Abstract

GCaMP-based calcium imaging is a powerful tool for investigating neural function in specific neurons. We generated transgenic (Tg) medaka strains expressing jGCaMP7s across extensive brain regions under the control of the *gap43* promoter. Using these Tg larvae, calcium imaging successfully detected a tricaine-induced suppression of spontaneous neural activity and topographical visual responses in the optic tectum elicited by moving paramecia or optical fiber stimulation. These results indicate that our Tg medaka strains provide a versatile platform for investigating neural dynamics and their responses to various stimuli.

## 1 Introduction

Calcium acts as a universal second messenger, regulating critical cellular signaling processes across diverse tissues and organisms (Tian et al., 2009). In neurons, action potentials trigger rapid intracellular calcium influx, while synaptic transmission via glutamate receptors induces Ca^2+^ transients in dendritic spines. Genetically encoded calcium indicators (GECIs), such as GCaMP (Nakai et al., 2001), are powerful tools for imaging neuronal activity and have been successfully applied in a wide range of species.

Zebrafish larvae, owing to their small size and transparency, are particularly well-suited for GCaMP-based calcium imaging, allowing visualization of neural activity from single neurons to whole-brain circuits (Vanwalleghem et al., 2018). Japanese medaka (*Oryzias latipes*) shares many advantages with zebrafish, such as embryo transparency (Furutani-Seiki and Wittbrodt, 2004), availability of genomic information (Ichikawa et al., 2017; Kasahara et al., 2007), and genome editing techniques (Ansai and Kinoshita, 2014; Watakabe et al., 2018). From an evolutionary perspective, zebrafish and medaka diverged from each other approximately 314-332 million years ago (Kasahara et al., 2007), resulting in different behavioral patterns, especially in reproductive behaviors. For example, while zebrafish rely on pheromones (olfactory signals) to trigger group spawning (Hutter et al., 2010; Yabuki et al., 2016), medaka possesses remarkable abilities in recognizing familiar individuals and selecting mates, primarily relying on visual information (Okuyama et al., 2014; Wang and Takeuchi, 2017; Yokoi et al., 2020). However, as calcium imaging techniques for the brain had not been established in medaka, we developed a whole-brain calcium imaging system using medaka larvae as our initial approach. Here, we generated transgenic medaka expressing jGCaMP7s (Dana et al., 2019) across broad brain regions and validated its use for calcium imaging to monitor neuronal activity in medaka.

## 2 Materials and Methods

### 2.1 Fish

The hi-medaka strain (MT835; https://shigen.nig.ac.jp/medaka/) provided from the National BioResource Project was used as the parental strain. All fish were bred and raised according to previously described protocols (Seki et al., 2023). Animal experiments were conducted with approval from Tohoku University’s animal care committee (permission number: 2022LsA-003).

### 2.2 Preparation of Ac/Ds transposon system

The construction of plasmid vectors and preparation of Ac nuclease is shown in the Supplemental Material and Methods.

### 2.3 Preparation of CRISPR/Cas system

Cas9 nuclease and sgRNA were prepared as previously described (Ansai and Kinoshita, 2014; Seki et al., 2023). The target sequences of sgRNAs used in this study were shown in Table S1.

### 2.4 Establishment of the transgenic strain

Injection mixtures containing Ac mRNA (25 ng/μl) and the donor plasmid pmDs-gap43:jGCaMP7s (10 ng/μl) (Figure S1) were injected into 1-cell stage medaka embryos. The injected fish with green fluorescent protein (GFP) expression in the brain were screenedusing an upright fluorescence stereo microscope (SZX16, Olympus). The GFP-positive fish were backcrossed with wild type (WT).

In addition, to eliminate the leucophores that interfere with calcium imaging by its autofluorescence, *slc2a15b*, which is essential for leucophore pigmentation in medaka (Kimura et al., 2014; Lischik et al., 2019), was disrupted by CRISPR/Cas system. Briefly, we injected Cas9 mRNA(100 ng/μl) and two sgRNAs for *slc2a15b* (25 ng/μl each) into1-cell stage embryos of the F_1_ generation of the transgenic strain. Then, fish with strong GFP fluorescence and minimal leucophore pigmentation were selected as founder fish. After crossing between the founder fish, the GCaMP-expressingtransgenic fish lacking leucophores Tg(gap43:jGCaMP7s)had been established. This generation and the subsequent generation were used for tricaine administration and various stimulus presentation experiments. For confocal microscopy, we used crispants of *slc45a2*, a gene involved in the synthesis of melanophores, that were generated by injection of Cas9 RNA and two sgRNAs (Table S1) as the *slc2a15b disruption*.

### 2.5 Fluorescence observations

For confocal microscopy, we used an A1R confocal system (Nikon) equipped with a 488 nm laser with an ApoLWD 25× objective lens, as previously reported (Hiyoshi et al., 2021). For wide-field imaging, we used a zoom microscope Axio Zoom.V16 (Zeiss) with a PlanNeoFluar Z 1.0× objective lens, green fluorescence protein (GFP) filter set 38 HE, RFP filter set 63 HE, a monochrome Axiocam 705 mono camera (Zeiss) and Zen Blue 3.7 software (Zeiss).

### 2.6 Wide-field Calcium imaging under tricaine treatment

We used 13 days post fertilization (dpf) Tg larvae for the experiment. After anesthetizing the larvae with tricaine (Sigma-Aldrich), they were embedded in 2% low-melting agarose (Agarose -LM, Nacalai Tesque) in a 55 mm dish. Small holes were created around the larvae with a φ0.1 m/m tungsten needle (Nisshin EM) to enhance drug permeability. After adding 3 ml of aquarium water, control imaging was performed twice at 4 Hz for 2 minutes. Subsequently, 1.5 ml of water was manually removed, and 1.5 ml of 1.2% tricaine solution (final concentration: 0.6%) was added. Imaging was carried out at 5 and 10 minutes post-administration, with focal adjustments made to match the control imaging conditions. Following imaging, the solution was replaced with aquarium water, and the larvae were released from the gel and allowed to recover from anesthesia. After a recovery period of at least one hour, the larvae were re-embedded in agarose, and two post-wash imaging sessions were conducted under the same conditions. All images were acquired under 100 × magnification with 4 × 4 binning at 608 × 514 pixels with a pixel size of 1.380 μm.

### 2.7 Wide-field Calcium imaging with visual stimuli

To eliminate pigmentated melanophores, we administered 0.003% PTU (N-phenylthiourea, Nacalai Tesque) to embryos from 2 dpf until hatching, and 3-7 days post-hatch Tg larvae were used for the experiment. The larvae were fed paramecium one time per day. For live paramecium presentation, the agarose around the head was removed, and 1-5 paramecia were introduced into the exposed space to swim freely. Imaging was conducted at 5 Hz. For optical fiber-based stimulation, the gel around the mouth and one eye was removed, and a manipulator moved the optical fiber connected to a red LED (OSR6LU5B64A-5V, OptoSupply). The on/off of the LED was controlled by a Raspberry Pi pico (Raspberry Pi Foundation). The optical fiber was moved around the exposed eye to deliver visual stimulation during imaging. Images were acquired under 50-63 × magnification with 4×4 or 5 × 5 binning to ensure that the movements of paramecium and fibers remained within the field of view.

### 2.8 Imaging data analysis

Image data were processed using Fiji of ImageJ (Schindelin et al., 2012). The acquired data of the tricaine treatment assay were first corrected for photobleaching using the Exponential Fit method in ImageJ Fiji’s bleach correction plugin (Miura, 2020). The average image of all analyzed frames was created as baseline (F_0_) and dF or dF/F was calculated (dF = F(t)-F_0_, dF/F_0_ = [F(t)-F_0_]/F_0_) for each movie data. In the tricaine treatment assay data, the region of interest (ROI) encompassing the entire brain was manually defined, and only the data within this ROI were extracted. Square ROIs with a side length of 30 pixels were arranged in a grid of 20 rows by 17 columns within a 608 x 514 pixel frame. Among them, only ROIs that were entirely contained within the extracted brain region were selected and the dF/F values were obtained. For peak detection, we referred to a previously reported method for whole-brain calcium imaging in zebrafish larvae (Turrini et al., 2017). First, filtering was performed using the Savitzky-Golay method with the savgol_filter function from Python’s SciPy library, with a polynomial order of 7 and a frame length of 25. Peaks were then automatically detected in the filtered data using the find_peaks function from the SciPy library, with parameters set to prominence = 0.2, height = 0, and distance = 25. Confocal imaging data and visual stimulus imaging data were subjected to 2×2 mean filtering to reduce noise.

### 2.9 Statistical analysis

All statistical analyses were performed using R version 4.2.0. The effects of tricaine administration on the number of peaks of dF/F were examined using the one-way aligned rank transform (ART) analysis of variance (ANOVA) in the R package ‘ARTool’ version 0.11.1. For the post hoc test, significant differences between all combinations of two factors were evaluated by Tukey’s multiple comparison test implemented in the ‘art.con’ function. The detailed statistical results are presented in Table S2.

## 3 Results

### 3.1 Generation of transgenic medaka expressing GCaMP in broad brain regions

To establish medaka expressing GECI in neuronal cells, we generated a transgenic strain expressing jGCaMP7s driven by medaka *gap43* promoter (Fujimori et al., 2008) using the Ac/Ds transposon system (Boon Ng and Gong, 2011; Froschauer et al., 2012)) (Figure 1A). To reduce leucophore autofluorescence interrupting fluorescence observation, we disrupted the slc2a15b gene using CRISPR/Cas system and then established a transgenic strain Tg(*gap43: jGCaMP7s*) without pigmented leucophores (*slc2a15b*^*-/-*^) (Figure 1B). The Tg larvae showed GFP fluorescence in the brain and spinal cord (Figure 1B). To examine whether the GCaMP-expressing cells are distributed widely in the brain at cellular resolution, we observed the brain using a confocal microscopy. The larvae showed GCaMP fluorescence signals throughout the brain (12-15 dpf, n = 3) (Figures 1C and 1D, Movie S1-S2) and part of the spinal cord (Figure 1B, white arrow). Prominent signals were observed in the habenula, nucleus pretectalis superficialis (PS) and the tectal longitudinal column (TL). These expression patterns were generally consistent with previous reports in GFP transgenic medaka driven by the *gap43* promoter (Fujimori et al., 2008; Kawasaki et al., 2023).

**Figure 1.**
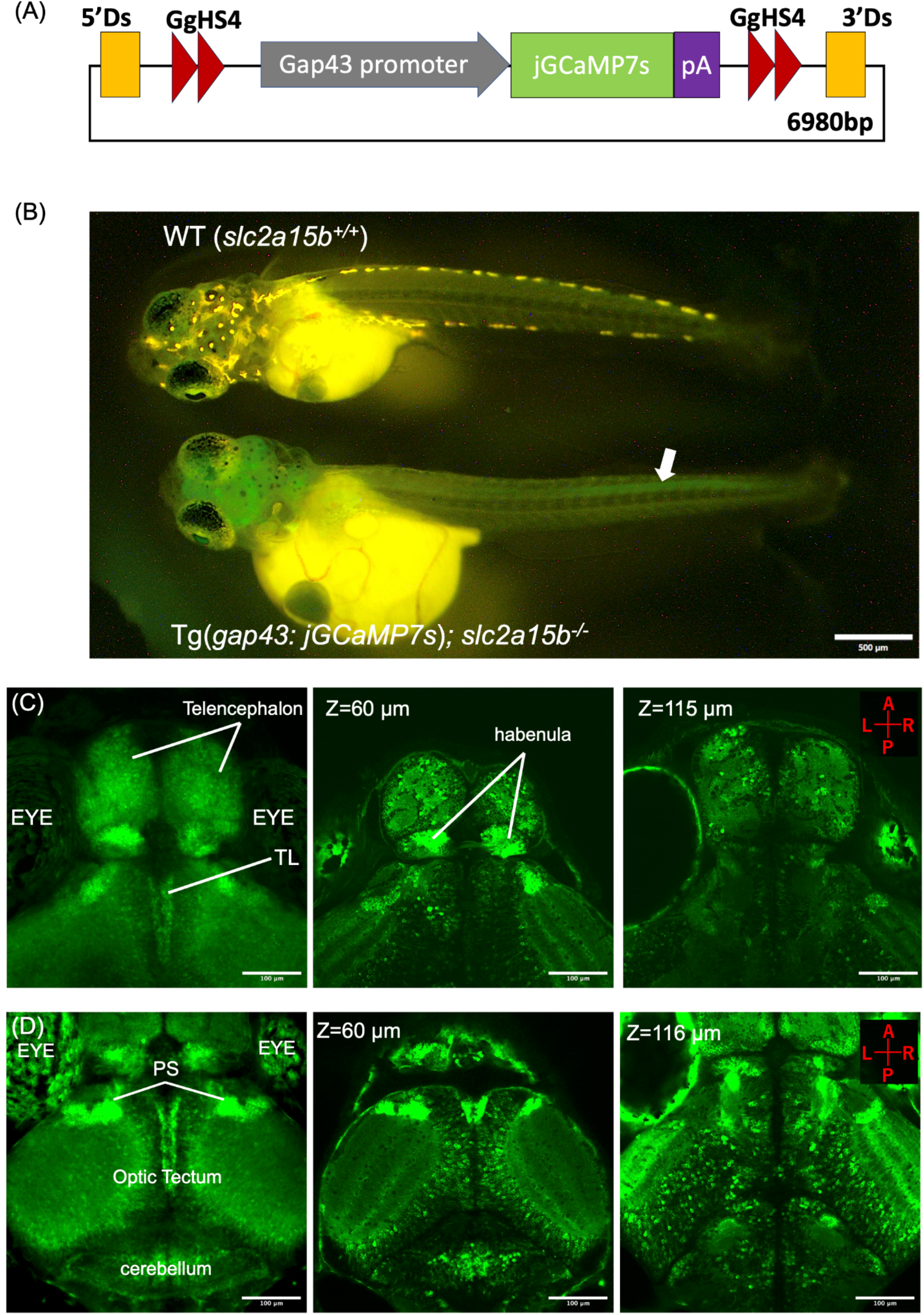
Generation of a transgenic medaka strain Tg(*gap43: jGCaMP7s*). (A) Schematic illustration of the transgenic construct. The construct contains jGCaMP7s gene with a SV40 polyA signal (pA) under the promoter derived from the Growth-associated protein 43 gene (gap43). The GCaMP expression cassette was separated from endogenous enhancer activity by inserting insulator sequences derived from chicken (GgHS4) (Shimizu and Shimizu, 2013). (B) Fluorescence image of WT and Tg(*gap43: jGCaMP7s*) larvae. In the WT larva, no GCaMP fluorescence was detected, while yellow autofluorescence from leucophores was visible. In contrast, Tg(gap43: *jGCaMP7s*) larva showed no leucophores, and autofluorescence was suppressed. GCaMP fluorescence was detected in both the brain and spinal cord. The white arrow indicates the GCaMP signal in the spinal cord. (C-D) Confocal microscopy of Tg(*gap43: jGCaMP7s*) expression pattern of jGaCaMP7s in the larval brain (10-13 dpf). (C-D) Left: Average intensity projection along the Z-axis of the telencephalon and rostral part of the optic tectum and cerebellum respectively. Middle and Right: Representative frame of Z-scan. Z represents the depth from the dorsal surface of the brain. A: anterior, P: posterior, R: right, L: left. PS: nucleus pretectalis superficialis, TL: torus longitudinalis.

### 3.2 GCaMP7s-based calcium imaging reveals spontaneous neural activity and their suppression by tricaine across the larval medaka brain

Next, we tested whether our transgenic medaka could detect spontaneous neural activity. Time-lapse imaging using both confocal and wide-field microscopy, revealed spontaneous fluctuations in fluorescence intensity across various brain regions in agarose-embedded transgenic larvae (Movies S3 and S4, respectively). To validate that these calcium signals represent neural activity, we performed calcium imaging during acute suppression of neuronal firing using tricaine (Figure 2). Tricaine is commonly used as an anesthetic for fish and amphibians, and electrophysiological studies have demonstrated that it primarily inhibits Na+ currents (Frazier and Narahashi, 1975; Wang et al., 1994) and suppresses neural activity in zebrafish (Svoboda et al., 2001). It also reduces calcium signals in whole-brain GCaMP imaging of zebrafish larvae (Turrini et al., 2017). Calcium imaging was performed in sequential phases: two baseline recordings prior to tricaine administration, two recordings during tricaine treatment (at 5 and 10 minutes post-administration), and then two recordings following tricaine washout and recovery from anesthesia (Figure 2A). To quantify neural activity levels, we counted the number of fluorescence intensity peaks. For systematic detection of localized neural activity, we employed automated grid-based ROI placement across the entire brain, followed by automated peak detection within each ROI (Figure 2B). Following the tricaine administration, the number of peaks of calcium transients decreased significantly; this effect was reversed upon tricaine washout, with the number of peak returning to baseline levels (one-way ART ANOVA, F = 13.9, df = 5, p < 0.001, Tukey’s post hoc test, p < 0.05; Figure 2C, Movie S5).

**Figure 2.**
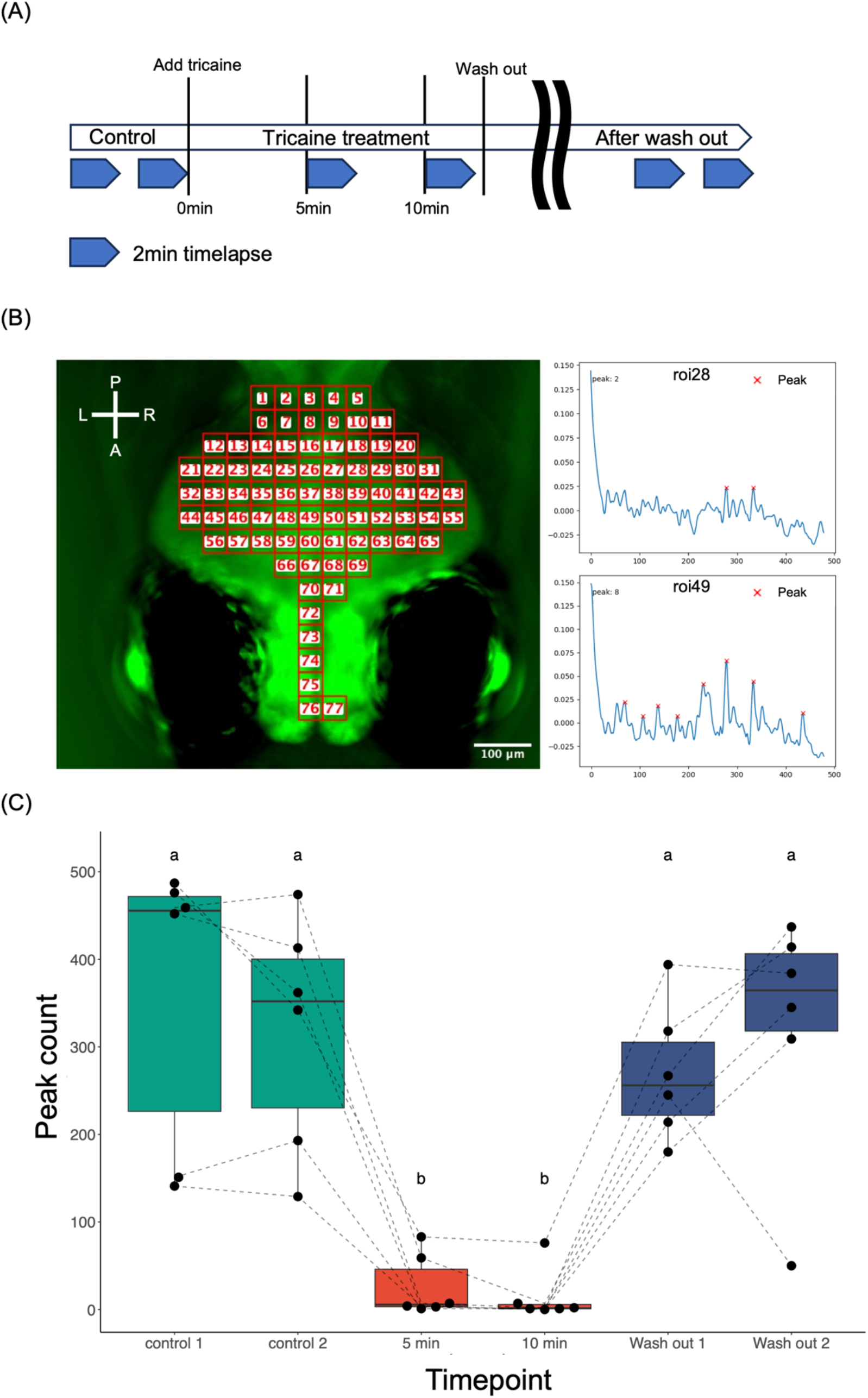
Effect of tricaine administration on spontaneous neural activity using wide-field microscopy. (A) Experimental timetable. (B) Left: An example of placing grid ROI over the entire brain. Right: Examples of automatic peak detection. The ROI number corresponds to the number of the grid ROI specified for the brain in the left panel. Red crosses indicate the positions of automatically detected peaks. (C) Box plot of the number of peaks at each time point. The boxes display the median, 25th, and 75th percentiles for each group. Each dot or line represents an individual. The lowercase letters above each box indicate the results of the two-way aligned rank transform (ART) analysis of variance (ANOVA) followed by post hoc Tukey’s multiple comparison tests and different letters indicate significant differences from each other (p < 0.05).

### 3.3 Visual stimuli from paramecia elicited topographically organized responses in the optic tectum

To monitor neural activities evoked by any stimuli using this transgenic fish, we conducted calcium imaging in medaka presenting two different visual stimuli (Figure 3). Previous studies have demonstrated the visuotopic organization of the larval zebrafish brain using GCaMP-based calcium imaging (Muto et al., 2013). Briefly, moving stimuli, such as swimming paramecia or computer-generated dots, evoked calcium responses in the optic tectum, that tracked the stimulus position, consistent with the retinotopic map previously established through anatomical studies (Baier et al., 1996; Stuermer, 1988) (Figure 3A). To investigate whether medaka fish exhibit similar responses, we recorded calcium dynamics while presenting live paramecia to agarose-embedded larvae. When calcium imaging was performed while presenting paramecia, responses were primarily observed in the optic tectum (Movie S6). Specifically, anteroposteriorly-directed paramecium movement elicited calcium transients in the anteroposterior regions of the optic tectum (Figure 3B, Movie S7) and dorsoventral-directed movement induced calcium transients in the ventral (central) to dorsal (marginal) regions of the optic tectum (Figure 3C, Movie S8). To further characterize the relationship between stimulus motion and calcium dynamics, we conducted controlled optical fiber stimulation. A light-emitting fiber was systematically moved along dorsoventrally and anteroposteriorly axes within the visual field (Figure 3D). Similar to the paramecium-evoked responses, optical fiber movement evoked calcium transient signals in the optic tectum. Anteroposterior-directed fiber movement induced anteroposterior calcium dynamics (Figure 3E, Movie S9) and dorsoventral-directed movement induced sequential calcium transients in ventral (central) to dorsal (marginal) regions (Figure 3F, Movie S10).

**Figure 3.**
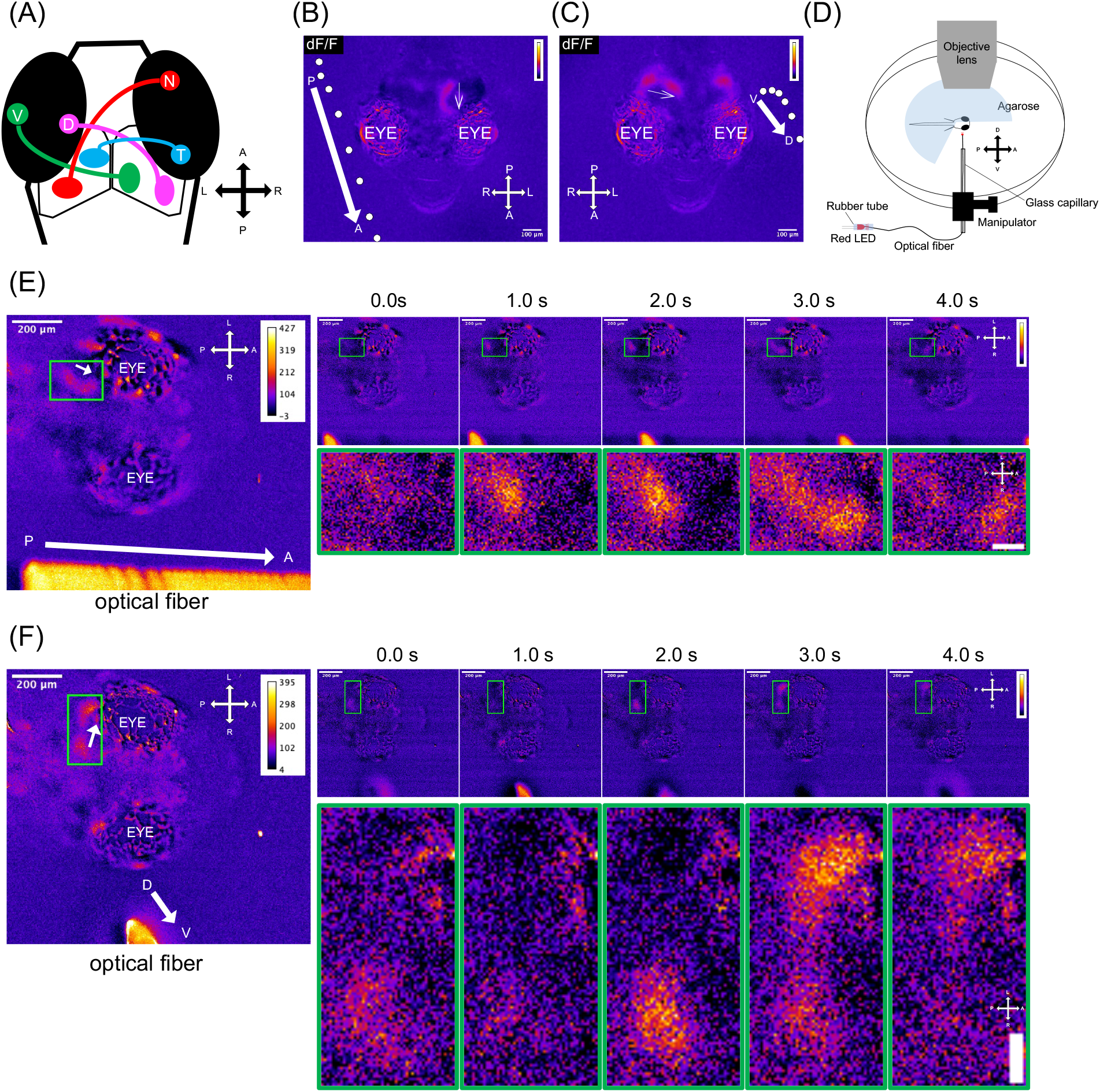
Neural responses in the optic tectum to visual stimuli. (A) Retinotopic projections of retinal ganglion cell axons. (B-C) Neural activity in response to a paramecium moving from anterior to posterior. Maximum intensity projection of ratio images (dF/F =[F(t)-F_0_]/F_0_) were pseudocolored. Calcium transients are indicated by a white thin arrow and the white dots and white thick arrow indicate the paramecium’s trajectory. (B) A paramecium moved from the posterior to the anterior, and calcium signals moved from the posterior to the anterior region in the optic tectum. (C) A paramecium moved from the ventral to the dorsal, calcium signals moved from the center to the margin region in the optic tectum. (D) Schematic of the optical fiber stimulation setup. (E-F) Frames showing calcium transients during optical fiber movement. Ratio images (dF = F(t)-F_0_) were pseudocolored. Left: Maximum intensity projections of the time-lapse. Right: Frame series showing fiber movement. (E) When the fiber moved from posterior to anterior, calcium transients moved in the left optic tectum from posterior to anterior. (F) When the fiber moved dorsoventrally, calcium transients shifted from the center to the margin of the left optic tectum. D, dorsal; V, ventral; N, nasal; T, temporal; A, anterior; P, posterior; L, left; R, right.

## 4 Discussion

In this study, we successfully implemented calcium imaging in larval medaka using a GECI to monitor neural activities. First, spontaneous fluctuations in GCaMP were observed in awake fish but not in fish treated with tricaine, indicating that the calcium signals accurately reflect the spontaneous neural activities in the larval medaka brain. In the visual stimulus experiments, both paramecium movement and optical fiber stimulation revealed a functional visuotopic map in the optic tectum in larval medaka. The topological relationships observed in these experiments were consistent with those reported in zebrafish (Muto et al., 2013). These findings suggest that our transgenic strain is well-suited for monitoring neural activity in response to various visual stimuli. To date, several transgenic medaka strains expressing GECIs have been reported; however, their expression has been restricted to the pituitary gland (Fukuda et al., 2025; Karigo et al., 2014). Thus, our strain established represents the first transgenic medaka strain exhibiting widespread expression of GECI throughout the brain.

Recent advances in targeted gene knock-in techniques in medaka (Kayo et al., 2024; Murakami et al., 2017; Seleit et al., 2021; Watakabe et al., 2018) have eliminated the labor-intensive process of promoter identification and cloning, thereby facilitating precise cell-type labeling. Furthermore, we validated the efficacy of binary expression systems, specifically the Tet system, for labeling specific neurons in both embryonic and adult medaka (Hosoya et al., 2021; Kayo et al., 2024). This system can be analogous to the GAL4/UAS enhancer/gene trap systems employed in zebrafish and is expected to accelerate the development of genetic tools for analyzing neurons and circuits genetically labeled by specific promoters and/or enhancers. Additionally, whole-brain imaging techniques have been established in zebrafish under free-swimming conditions (Hasani et al., 2023; Kim et al., 2017) and in more complex behavioral contexts (Migault et al., 2018). Recent studies have also demonstrated the feasibility of imaging adult zebrafish (Huang et al., 2020; Torigoe et al., 2021). Combining calcium imaging with these advanced genetic tools and imaging techniques is expected to enable the functional characterization of neurons in medaka, particularly those involved in visual processing and mate choice behavior.

## Acknowledgements

We thank the National BioResource Project Medaka (https://shigen.nig.ac.jp/medaka) for providing the hi-medaka strain (ID: MT835). We also thank Kazuhide Miyamoto and Prof. Koji Tamura of the Laboratory of Organ Morphogenesis for providing live paramecium. Finally, we thank laboratory members of Molecular Ethology for valuable discussions and technical support.

