## Supplemental Material and Methods for "Establishment of a transgenic strain for the whole brain calcium imaging in larval medaka fish (*Oryzias latipes*)"

### Supplemental information

#### Supplemental Materials and Methods

##### *Ac mRNA preparation*

Capped mRNA for Ac transposase was synthesized as described previously (Ishikawa et al., 2018) with slight modifications. Briefly, the expression vector pCS2+Ac was linearized with the NotI digestion and then purified with proteinase K treatment followed by spin column purification using NucleoSpin Gel and PCR Clean-up kit (Macherey–Nagel) with Buffer NTB. The capped mRNA for Ac transposase was synthesized using mMessage mMachine SP6 kit (Invitrogen) and then purified using Monarch RNA Cleanup kit (50 µg) (NEB).

##### *Plasmid vector construction*

For the transgenic construct pmDs-gap43-jGCaMP7s, the jGCaMP7s gene was amplified from the vector pGP-CMV-jGCaMP7s (Addgene plasmid #104463) (Dana et al., 2019). A 2.0-kb of medaka *gap43* promoter (Fujimori et al., 2008) was amplified from medaka genomic DNA using primers gap43-proFW (5'-aaaCTCGAGttttcacaccatcctttgagg-3') and gap43-proRV (5'-ggcACTAGTagttggacttgtagtatcctcctgt-3'). The PCR fragments were cloned with an SV40 polyA signal into pmDs vector containing 5′- and 3′-Ds sequences and two tandem repeats of chicken-derived insulator (cHS4) (Ishikawa et al., 2018). The full sequence of the transgenic construct is available in the GitHub repository (https://github.com/satoshi-ansai/Plasmids).

Table S1: List of the target sequences of sgRNAs used in this study.

| Name | Target sequence (5’ – 3’) |
|---|---|
| T7-sgRNA-slc2a15b#1 | CAGGTGAGTTGACGACTGCCAGG |
| T7-sgRNA-slc2a15b#2 | ATGAACGGATATGGAACTCTTGG |
| T7-sgRNA-slc45a2#1 | ACGGAGCGCACTGGAAGCTGAGG |
| T7-sgRNA-slc45a2#1 | CCAAGGTCCTACTCGGCGATTGG |

Table S2: Summary of the results of transgenics created in this study.

one-way aligned rank transform (ART) analysis of variance (ANOVA)

| Term | Df | Df.res | Sum Sq | Sum Sq.res | F value | Pr(>F) |
|---|---|---|---|---|---|---|
| timepoint | 5 | 30 | 2695.75 | 1160.75 | 13.93453 | 4.52E-07 |

post hoc test with Tukey's multiple comparison

| Group1 | Gruop2 | estimate | SE | df | t.ratio | p.value |
|---|---|---|---|---|---|---|
| control1 | control2 | -0.166667 | 3.59127 | 30 | -0.046409 | 1 |
| control1 | tri5 | 17.75 | 3.59127 | 30 | 4.942541 | 0.000364 |
| control1 | tri10 | 21.25 | 3.59127 | 30 | 5.917126 | 2.43E-05 |
| control1 | Wash out1 | 7.583333 | 3.59127 | 30 | 2.111602 | 0.308627 |
| control1 | Wash out2 | 1.083333 | 3.59127 | 30 | 0.301657 | 0.999628 |
| control2 | tri5 | 17.91667 | 3.59127 | 30 | 4.98895 | 0.00032 |
| control2 | tri10 | 21.41667 | 3.59127 | 30 | 5.963535 | 2.14E-05 |
| control2 | Wash out1 | 7.75 | 3.59127 | 30 | 2.158011 | 0.286516 |
| control2 | Wash out2 | 1.25 | 3.59127 | 30 | 0.348066 | 0.999254 |
| tri5 | tri10 | 3.5 | 3.59127 | 30 | 0.974586 | 0.922428 |
| tri5 | Wash out1 | -10.16667 | 3.59127 | 30 | -2.830939 | 0.079627 |
| tri5 | Wash out2 | -16.66667 | 3.59127 | 30 | -4.640883 | 0.000834 |
| tri10 | Wash out1 | -13.66667 | 3.59127 | 30 | -3.805524 | 0.007759 |
| tri10 | Wash out2 | -20.16667 | 3.59127 | 30 | -5.615469 | 5.62E-05 |
| Wash out1 | Wash out2 | -6.5 | 3.59127 | 30 | -1.809945 | 0.474829 |

### Legends to Supplemental Movies

**Movie S1.** Telencephalon and optic tectum region of horizontal optical sections showing GCaMP expression pattern in 13 dpf larva of *slc45a2* crispant of Tg(*gap43: jGCaMP7s/ slc2a15*$^{-/-}$). Video displays the z-stack moving from dorsal to ventral.

**Movie S2.** Optic tectum and cerebellum region of horizontal optical sections recorded under conditions similar to those used in Movie S1.

**Movie S3.** Representative results of spontaneous calcium transients observed with time-lapse confocal imaging. The left panel displays the raw fluorescence intensity, while the right movie presents pseudocolored images of dF/F. The movie is played at 8× the real-time speed.

**Movie S4.** Representative results of spontaneous calcium transients observed with time-lapse wide-field imaging. The movie presents pseudocolored images of dF/F. The movie is played at 5× the real-time speed.

**Movie S5.** Tricaine administration experiments on a single Tg(*gap43: jGCaMP7s/ slc2a15*$^{-/-}$) larva. The four videos at each time point are displayed. The movie is played at 5× the real-time speed.

**Movie S6.** Representative calcium imaging during the presentation of several paramecia. Entry of the paramecium into the field of view induces calcium transients and travels primarily within the optic tectum. The movie is played at 6×the real-time speed.

**Movie S7.** Representative jGCaMP7s optical measurements during anteroposterior movement of the paramecium. When paramecia moved from the anterior to posterior area in the left visual field, calcium signals moved from anterior to posterior in the right optic tectum. When the paramecium moved from the posterior to the anterior area in the left visual field, calcium signals moved from center to margin in the right optic tectum. The movie is played at 1× the real-time speed.

**Movie S8.** Representative jGCaMP7s optical measurements during dorsoventral movement of the paramecium. When paramecia moved from the ventral to the dorsal area in the left visual field, calcium signals moved from the margin to the center in the right optic tectum. When the paramecium moved from the dorsal to the ventral area in the right visual field, calcium signals moved from posterior to anterior in the left optic tectum. The movie is played at 1× the real-time speed.

**Movie S9.** Representative jGCaMP7s optical measurements during stimulation with the anteroposterior movement of the optical fiber. A transgenic larva was embedded in agarose, and an optical fiber was moved in front of the larva's right eye. In the area indicated by the red arrow, the signal dynamics correspond to the anteroposterior functional visuotopic map in the optic tectum. The movie is played at 5× the real-time speed.

**Movie S10.** Representative results of jGCaMP7s optical measurements during stimulation with the dorsoventral movement of the optical fiber. The recording conditions are the same as those used in Movie S6. In the area indicated by the red arrow, the signal dynamics correspond to the dorsoventral functional visuotopy in the optic tectum. "D→V" or "V→D" displayed in the movie indicates the direction of the optical fiber movement. The movie is played at 5× the real-time speed. D: dorsal; V: ventral.
